# Protein engineering expands the effector recognition profile of a rice NLR immune receptor

**DOI:** 10.1101/611152

**Authors:** JC De la Concepcion, M Franceschetti, R Terauchi, S Kamoun, MJ Banfield

## Abstract

Plant NLR receptors detect pathogen effectors and initiate an immune response. Since their discovery, NLRs have been the focus of protein engineering to improve disease resistance. However, this has proven challenging, in part due to their narrow response specificity. Here, we used structure-guided engineering to expand the response profile of the rice NLR Pikp to variants of the rice blast pathogen effector AVR-Pik. A mutation located within an effector binding interface of the integrated Pikp-HMA domain increased the binding affinity for AVR-Pik variants in vitro and in vivo. This translates to an expanded cell death response to AVR-Pik variants previously unrecognized by Pikp in planta. Structures of the engineered Pikp-HMA in complex with AVR-Pik variants revealed the mechanism of expanded recognition. These results provide a proof-of-concept that protein engineering can improve the utility of plant NLR receptors where direct interaction between effectors and NLRs is established, particularly via integrated domains.

## Introduction

Protein engineering offers opportunities to develop new or improved molecular recognition capabilities that have applications in basic research, health and agricultural settings. Protein resurfacing, where the properties of solvent-exposed regions are changed (often by mutation), has been used extensively in diverse areas from antibody engineering for clinical use, to production of more stable, soluble proteins for biotechnology applications (1).

Intracellular nucleotide binding, leucine rich repeat (NLR) receptors are key components of plant innate immunity pathways. They recognise the presence or activity of virulence-associated, host-translocated pathogen effector proteins and initiate an immune response (2, 3). As they confer resistance to disease, plant NLRs are widely used in crop breeding programs (4). However, the recognition spectrum of plant NLRs tends to be very specific, and pathogens may delete detected effectors from their genome or evolve novel effector variants not detected by NLRs to reestablish disease (5).

The potential of engineering NLRs to overcome these limitations, or to detect new effector activities, is emerging (6). Some success has been achieved in expanding the activation sensitivity or effector recognition profiles of NLRs through gain-of-function random mutagenesis (7–9). In an alternative strategy, NLR perception of protease effectors, through their activity on engineered host proteins, can lead to expanded recognition profiles (10–12). When detailed knowledge of direct binding interfaces between an effector and an NLR are known, this offers the potential for protein resurfacing to modify interactions, and impact immune signalling.

Plant NLRs are modular proteins, defined by their nucleotide-binding (NB-ARC) and leucine rich repeat (LRR) domains, but also have either an N-terminal coiled-coil (CC) or Toll/Interleukin-1/Resistance-protein (TIR) signalling domain (13). However, many NLRs also contain non-canonical integrated domains (14–16). Integrated domains are thought to be derived from ancestral virulence-associated effector targets, which directly bind pathogen effectors (or host proteins (17)) to initiate an immune response (18–22). As such, these domains present an exciting target for protein engineering approaches to improve NLR activities. NLRs containing integrated domains (often called the “sensor”) typically function in pairs, requiring a second genetically linked NLR (the “helper”) for immune signalling (23,24).

Two rice NLR pairs, Pik and Pia, contain an integrated Heavy Metal Associated (HMA) domain in their sensor NLR that directly binds effectors from the rice blast pathogen *Magnaporthe oryzae* (also known as *Pycularia oryzae*) (20,25–27). The integrated HMA domain in the sensor NLR Pik-1 directly binds the effector AVR-Pik. Co-evolutionary dynamics has driven the emergence of polymorphic Pik-1 HMA domains and AVR-Pik effectors in natural populations, and these display differential disease resistance phenotypes (28–30). The Pikp NLR allele only responds to the effector variant AVR-PikD, but the Pikm allele responds to AVR-PikD, AVR-PikE, and AVR-PikA. These phenotypes can be recapitulated in the model plant *Nicotiana benthamiana* using a cell death assay, and are underpinned by differences in effector/receptor binding interfaces that lead to different affinities in vitro (20,22).

We hypothesised that by combining naturally occurring favourable interactions observed across different interfaces, as defined in different Pik-HMA/AVR-Pik structures (20,22), we could generate a Pik NLR with improved recognition profiles. Here, we graft an interface from Pikm onto Pikp by mutating two residues in Pikp (Asn261Lys, Lys262Glu), forming Pikp^NK-KE^. This single-site mutation strengthens the cell death response in *N. benthamiana* to AVR-PikD, and gains a Pikm-like response to AVR-PikE and AVR-PikA. We show that this gain-of-function phenotype correlates with increased binding affinity of the effectors by the Pikp^NK-KE^-HMA domain in vitro and in vivo, and demonstrate this mutation results in a Pikm-like structure for Pikp^NK-KE^, when in complex with AVR-Pik effectors. Finally, we confirm that the newly engineered interface is responsible for the expanded response of Pikp^NK-KE^ by mutation of the effectors.

This study serves as a proof-of-concept for the use of protein resurfacing by targeted mutation to develop plant NLR immune receptors with new capabilities. In the future, such approaches have the potential to improve disease resistance in crops.

## Results

### Structure-informed engineering expands Pikp-mediated effector recognition in *N. benthamiana*

By comparing protein interfaces in the structures of Pikp-HMA and Pikm-HMA bound to different AVR-Pik effectors (20,22), we hypothesised that we could engineer expanded effector recognition capabilities to Pikp by point mutation. We constructed a series of mutations in the previously identified interface 2 and interface 3 regions of Pik-HMA/AVR-Pik structures (22), swapping residues found in Pikm into Pikp (**Figure 1A,Figure 1 - figure supplement 1A**). We then screened these mutations for expanded effector recognition by monitoring cell death in a well-established *N. benthamiana* assay (20,22). We found one double mutation in two adjacent amino acid residues contained within interface 3, Asn261Lys and Lys262Glu (henceforth Pikp^NK-KE^), showed cell death in response to AVR-PikE and AVR-PikA (**Figure 1,Figure 1 - figure supplement 1A**). All proteins were confirmed to be expressed in plants by western blot (**Figure 1 - figure supplement 1B**).

**Figure 1.**
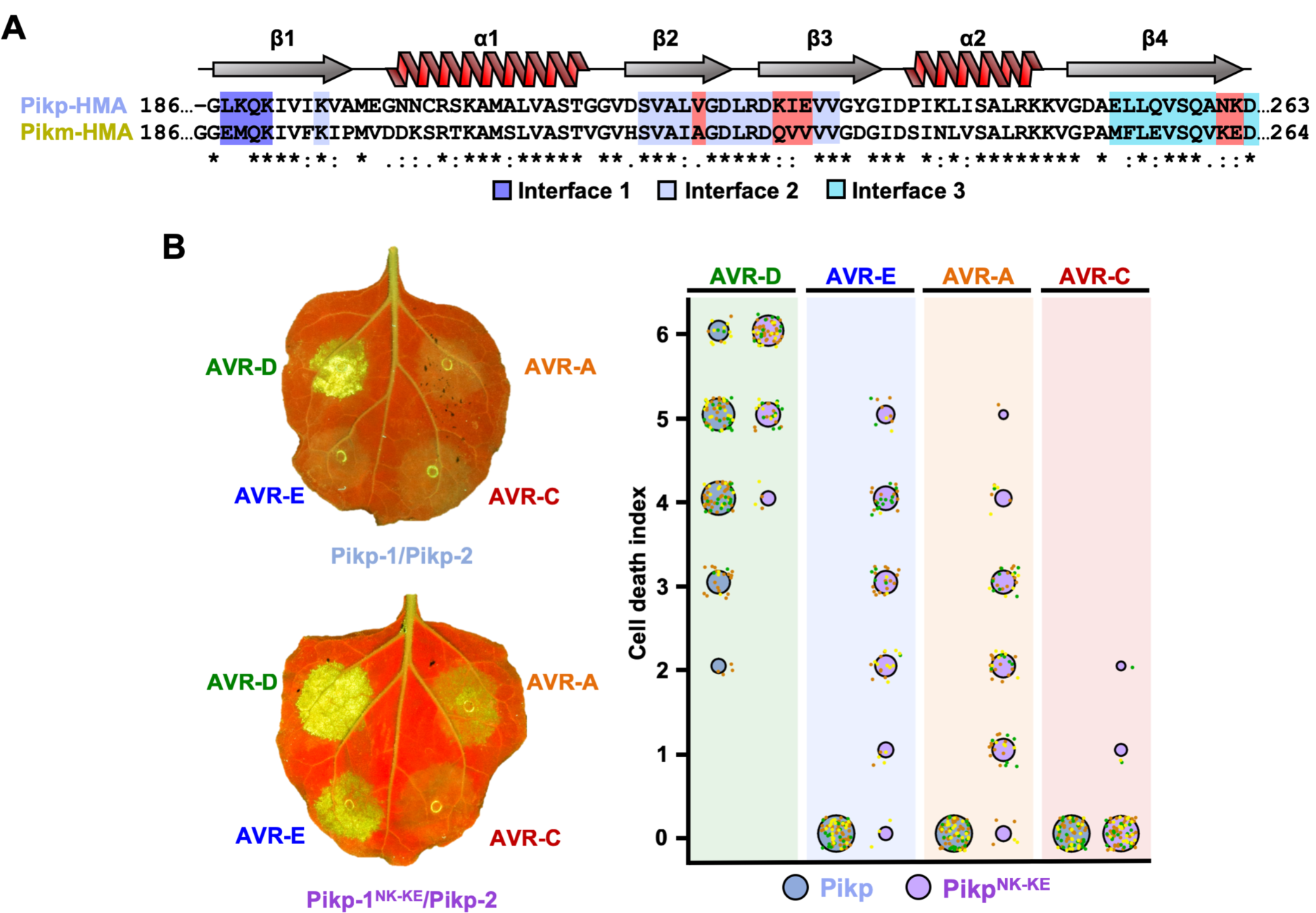
Structure-informed engineering expands Pikp-mediated effector recognition in *N. benthamiana*. **(A)** Sequence alignment of Pikp-1 and Pikm-1 HMA domains. Secondary structure features of the HMA fold are shown above, and the residues located to binding interfaces are as coloured. Key residues from interface 2 and interface 3 involved in this study are highlighted in red. **(B)** Left - A representative leaf image showing Pikp or Pikp-1^NK-KE^-mediated cell death to AVR-Pik variants as autofluorescence under UV light. Right - autofluorescence intensity is scored as previously described (20,22). Cell death assay scoring is represented as dot plots for Pikp and Pikp^NK-KE^ (blue and purple respectively). For each sample, all the data points are represented as dots with a distinct colour for each of the three biological replicates; these dots are jittered about the cell death score for visualisation purposes. The size of the centre dot at each cell death value is directly proportional to the number of replicates in the sample with that score. The total number of repeats was 80. Data for Pikp has been previously shown (22), but was acquired at the same time as Pikp^NK-KE^.

We subsequently focussed on this mutant, and independently repeated the cell death assay to ensure its robustness (**Figure 1B**). Similar to the Pikm allele (22), we observe a hierarchy of AVR-PikD>AVR-PikE>AVR-PikA for the intensity of cell death mediated by Pikp^NK-KE^, although in each case Pikp^NK-KE^ shows a qualitatively stronger response compared to Pikm (**Figure 1,figure supplement 2**). This is also observed when comparing the cell death of Pikp^NK-KE^ with Pikp in response to AVR-PikD (**Figure 1B**). Pikp^NK-KE^ does not show a response to the stealthy AVR-PikC variant in this assay.

We conclude that the single Asn261Lys/Lys262Glu (Pikp^NK-KE^) mutation at interface 3 in the Pikp NLR expands this protein’s recognition profile towards effector variants AVR-PikE and AVR-PikA, similar to that observed for Pikm.

### The engineered Pikp^NK-KE^-HMA mutant shows increased binding to effector variants in vivo and in vitro

We used Yeast-2-hybrid (Y2H) and surface plasmon resonance (SPR) to determine whether the expanded Pikp^NK-KE^ cell death response in *N. benthamiana* correlates with increased binding affinity of the Pikp^NK-KE^-HMA domain for AVR-Pik effectors.

As AVR-PikE and AVR-PikA show some interaction with Pikp-HMA using these approaches (20,22), we tested interactions with Pikp^NK-KE^-HMA side-by-side with wild-type. By Y2H we observed a small increase in growth/blue colouration (both indicative of protein-protein interaction) for Pikp-HMA^NK-KE^ with effectors AVR-PikE and AVR-PikA when compared with Pikp-HMA (**Figure 2A**). Unexpectedly, we observed some yeast growth for Pikp-HMA^NK-KE^ with AVR-PikC, comparable to Pikp-HMA with AVR-PikA (**Figure 2A**). Expression of all proteins was confirmed in yeast (**Figure 2 - figure supplement 1**).

**Figure 2.**
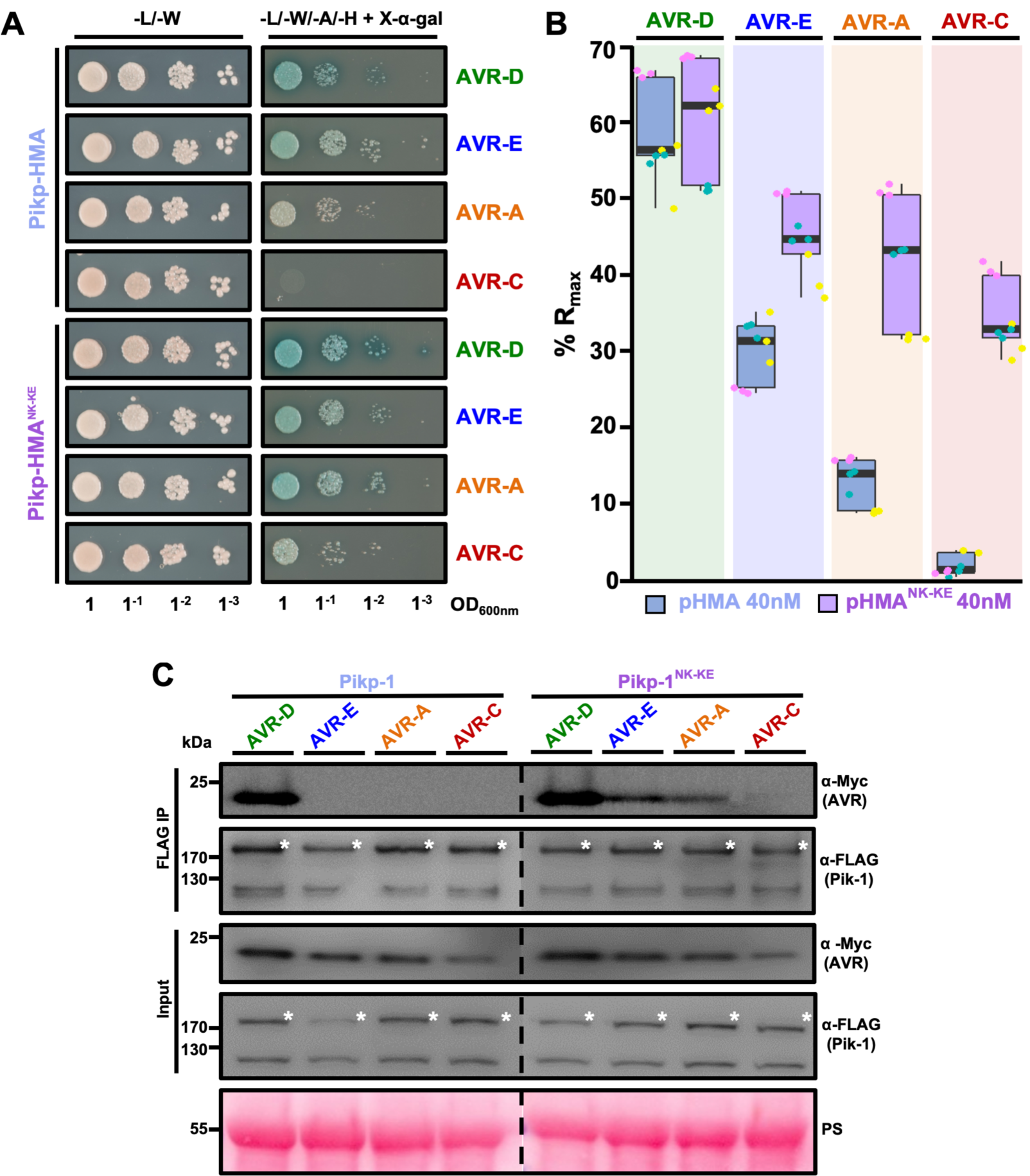
Pikp^NK-KE^ shows increased binding to effector variants in vivo and in vitro compared to wild type. **(A)** Yeast-Two-Hybrid assay of Pikp-HMA and Pikp-HMA^NK-KE^ with AVR-Pik alleles. Control plate for yeast growth is on the left, with the selective plate on the right for each combination of HMA/AVR-Pik. Growth and development of blue colouration in the selection plate are both indicative of protein:protein interaction. Each experiment was repeated a minimum of three times, with similar results. **(B)** Box plots showing %R_max_ for Pikp-HMA and Pikp-HMA^NK-KE^ with the AVR-Pik effectors alleles at HMA concentration of 40 nM as measured by surface plasmon resonance. Pikp-HMA and Pikp-HMA^NK-KE^ are represented by blue and purple boxes, respectively. The centre line represents the median, the box limits are the upper and lower quartiles, the whiskers are the 1.5× interquartile range and all the data points are represented as dots with distinct colours for each biological replicate. For each experiment, three biological replicates with three internal repeats were performed. For results of experiments with 4 and 100 nM HMA protein concentration see **Figure 2 - figure supplement 2**. **(C)** Co-immunoprecipitation of full length Pikp-1 and Pikp-1^NK-KE^ with AVR-Pik variants. N-terminally 4xMyc tagged AVR-Pik effectors were transiently co-expressed with Pikp-1:6xHis3xFLAG (left) or Pikp-1^NK-KE^:6xHis3xFLAG (right) in *N. benthamiana*. Immunoprecipitates (IPs) obtained with anti-FLAG antiserum, and total protein extracts, were probed with appropriate antisera. Dashed line indicates a crop site on the same blot used to compose the figure. Each experiment was repeated at least three times, with similar results. The asterisks mark the Pik-1 band, PS = Ponceau Stain.

Then, we produced the Pikp-HMA^NK-KE^ domain protein via overexpression in *E. coli* and purified it to homogeneity using well-established procedures for these domains (see **Materials and Methods**, (20,22)). Using SPR, we measured the binding affinity of the Pikp-HMA^NK-KE^ domain to AVR-Pik effectors, alongside wild-type Pikp-HMA, and also Pikm-HMA (**Figure 2b, Figure 2 - figure supplement 2,3**). Response units (RU) were measured following injection of Pik-HMAs at three different concentrations, after capturing AVR-Pik effectors on a Biacore NTA chip. RUs were then normalised to the theoretical maximum response (R_max_), assuming a 2:1 interaction model for Pikp-HMA and Pikp-HMA^NK-KE^, and 1:1 for Pikm-HMA, as previously described (22). This data showed an increased binding of Pikp-HMA^NK-KE^ to all AVR-Pik effectors compared to wild-type (**Figure 2b, Figure 2 - figure supplement 2**). The binding of Pikp-HMA^NK-KE^ to the AVR-Pik effectors was also consistently higher compared to Pikm-HMA (**Figure 2 - figure supplement 3**), correlating with cell death assays (**Figure 1 - figure supplement 2**). Although neither Pikp-HMA nor Pikm-HMA domains show binding to AVR-PikC by SPR, we observe a gain-of-binding of this effector variant with Pikp-HMA^NK-KE^ (**Figure 2b, Figure 2 - figure supplement 2, 3**), similar to the Y2H result.

These results show that the Pikp-HMA^NK-KE^ mutant has a higher binding affinity for effectors AVR-PikE and AVR-PikA than wild-type protein. This suggests that the increased binding affinity to the HMA domain correlates with the expanded cell death response in planta (**Figure 1B**).

### The engineered Pikp^NK-KE^ mutant expands association of full-length Pik-1 to effector variants in planta

In addition to interaction with the isolated HMA domain, we tested whether the Asn261Lys/Lys262Glu mutant could expand effector variant binding in the context of the full-length NLR. After generating the mutant in the full-length protein, we co-expressed either Pikp-1 or Pikp-1^NK-KE^ with the AVR-Pik effector variants in *N. benthamiana*, followed by immunoprecipitation and western blotting to determine effector association.

AVR-PikD shows a robust interaction with Pikp-1. However, although we observe limited binding for the isolated Pikp-HMA domain by Y2H and SPR, we did not detect association of AVR-PikE and AVR-PikA with the full-length Pikp-1 in planta (**Figure 2C**). By contrast, we observe clear association of AVR-PikE and AVR-PikA with the Pikp-1^NK-KE^ mutant, albeit with reduced intensity compared to AVR-PikD, correlating with the hierarchical cell death response observed in planta (**Figure 1C**).

We also detect a very low level of interaction between full length Pikp-1^NK-KE^ and AVR-PikC (**Figure 2C**). However, co-expression of Pikp-1^NK-KE^ and AVR-PikC does not result in macroscopic cell death in *N. benthamiana* (**Figure 1C**).

These results show that effector variant binding to full-length Pikp-1 and Pikp-1^NK-KE^ correlates with the in planta cell death response (**Figure 1C**).

### The effector-binding interface in the Pikp^NK-KE^ mutant adopts a Pikm-like conformation

Having established that the Pikp^NK-KE^ mutant displays an expanded effector recognition profile compared to wild-type, we sort to determine the structural basis of this activity. To this end, we determined crystal structures of Pikp-HMA^NK-KE^ bound to AVR-PikD, and to AVR-PikE. We obtained samples of Pikp-HMA^NK-KE^/AVR-PikD and Pikp-HMA^NK-KE^/AVR-PikE complexes by co-expression in *E. coli* (described in the **Materials and Methods** and (22)). Each complex was crystallised (see **Materials and Methods**) and X-ray diffraction data were collected at the Diamond Light Source (Oxford, UK) to 1.6 Å and 1.85 Å resolution respectively. The details of X-ray data collection, structure solution, and completion are given in the **Materials and Methods** and **Table 1**.

**Table 1:** Data Collection and Refinement statistics

|  | Pikp<sup>NK-KE</sup>:AVR-PikD | Pikp<sup>NK-KE</sup>:AVR-PikE |
| --- | --- | --- |
| **Data collection statistics** |  |  |
| Wavelength (Å) | 0.9763 | 0.9763 |
| Space group | *P* 2<sub>1</sub> 2<sub>1</sub> 2<sub>1</sub> | *P* 2<sub>1</sub> 2<sub>1</sub> 2<sub>1</sub> |
| Cell dimensions |  |  |
| *a*, *b*, *c* (Å) | 29.79, 65.33, 75.86 | 66.46, 80.70, 105.58 |
| Resolution (Å)* | 32.80-1.60 (1.63-1.60) | 29.50-1.85 (1.89-1.85) |
| *R*<sub>merge</sub> (%)# | 8.1 (97.1) | 5.2 (75.1) |
| *I*/σ*I*# | 16.1 (2.6) | 31.0 (4.1) |
| Completeness (%)# | 100 (100) | 99.8 (97.8) |
| Unique reflections# | 20294 (978) | 49337 (2963) |
| Redundancy# | 12.8 (13.3) | 18.3 (17.8) |
| CC<sup>(1/2)</sup> (%)# | 99.9 (86.6) | 100 (95.2) |
| **Refinement and model statistics** |  |  |
| Resolution (Å) | 32.82-1.60 (1.64 – 1.60) | 29.52-1.85 (1.90 – 1.85) |
| *R*<sub>work</sub>/*R*<sub>free</sub> (%)^ | 19.7/23.2 (25.5/27.3) | 18.6/23.0 (29.1/35.0) |
| No. atoms (Protein) | 1277 | 3604 |
| B-factors (Protein) | 25.6 | 39.7 |
| R.m.s deviations^ |  |  |
| Bond lengths (Å) | 0.009 | 0.012 |
| Bond angles (°) | 1.5 | 1.4 |
| Ramachandran plot (%)** |  |  |
| Favoured | 98.1 | 97.3 |
| Outliers | 0 | 0.2 |
| MolProbity Score | 1.41 (93<sup>th</sup> percentile) | 1.59 (91<sup>st</sup> percentile) |

\*The highest resolution shell is shown in parenthesis.

The overall architecture of these complexes is the same as observed for all Pik-HMA/AVR-Pik effector structures (20, 22), and an analysis of interface properties is given in **Figure 3 - figure supplement 1**. A key interaction at interface 3, one of the previously defined Pik-HMA/AVR-Pik interfaces (22), involves a Lysine residue (Lys262 in Pikp and Pikm) that forms intimate contacts within a pocket on the effector surface (**Figure 3**). In order to position this Lysine in the effector pocket, Pikp has to loop-out regions adjacent to this residue, compromising the packing at the interface ((22), **Figure 3A (left panel), B (left panel), C and D**). By contrast, in Pikm, where the position of the Lysine is one residue to the N-terminus, no looping-out is required to locate the Lysine into the pocket (**Figure 3A (right panel), B (right panel), C and D**). In the Pikp^NK-KE^ mutant, the position of this key Lysine is shifted one residue to the N-terminus compared to wild-type, and occupies the same position in the sequence as in Pikm. In the crystal structures of Pikp-HMA^NK-KE^ in complex with either AVR-PikD or AVR-PikE, we see that this region of the HMA adopts a Pikm-like conformation (**Figure 3A (middle panel), B (middle panel), C and D**), with no looping-out of the preceding structure. This confirms that with the Pikp^NK-KE^ mutant we have resurfaced Pikp to have a more robust, Pikm-like interface in this region.

**Figure 3.**
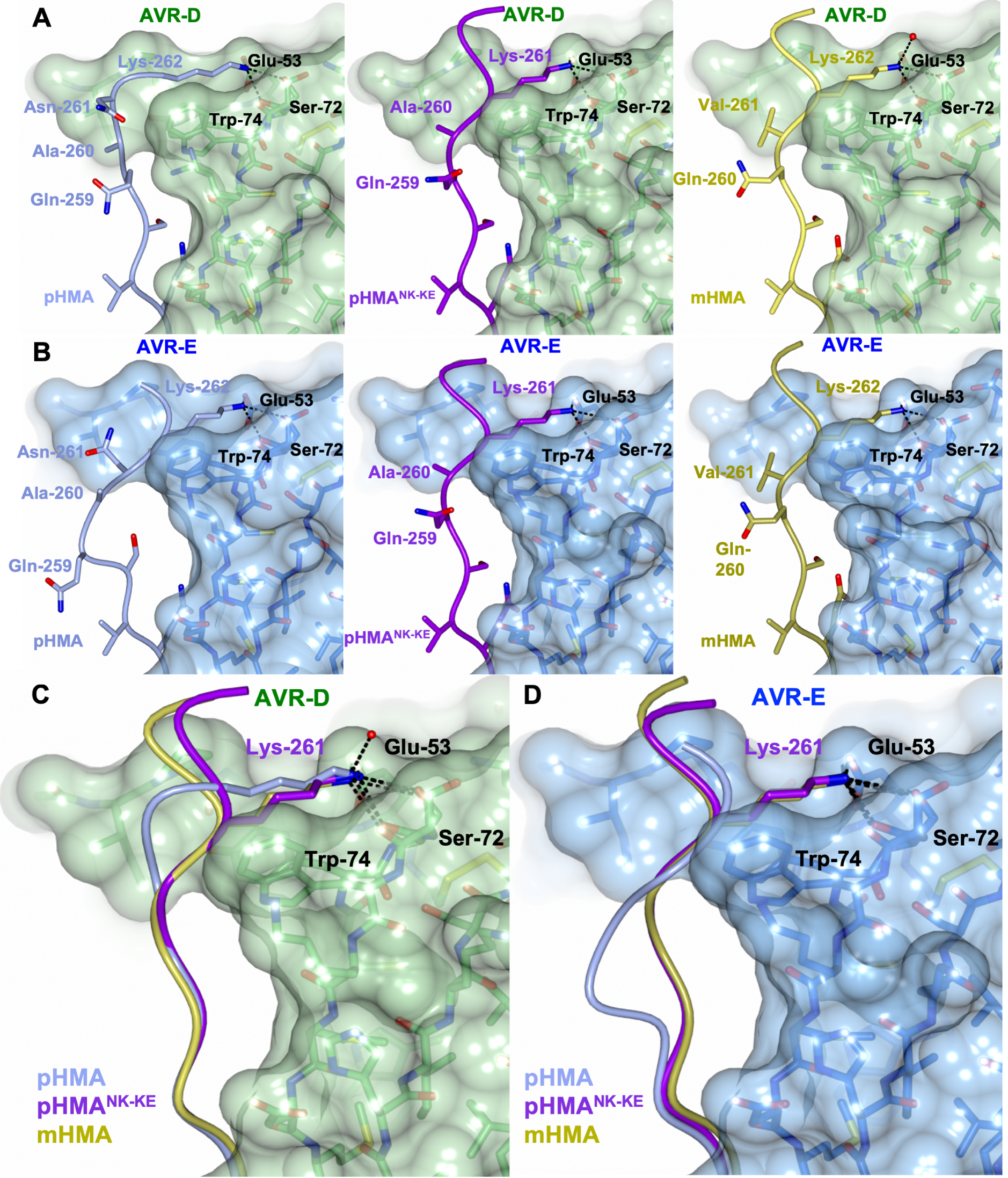
The Pikp^NK-KE^-HMA mutant adopts a Pikm-like conformation at the effector binding interface. Schematic view of the different conformations adopted by Pikp-HMA, Pikp-HMA^NK-KE^ and Pikm-HMA at interface 3 in complex with AVR-PikD or AVR-PikE. In each panel, the effector is shown as sticks with the molecular surface also shown and coloured as labelled. Pik-HMA residues are coloured as labelled and shown in the Cα-worm with side-chain representation. **(A)** Schematic of Pikp-HMA (left), Pikp-HMA^NK-KE^(middle) and Pikm-HMA (right) bound to AVR-PikD. Important residues involved in HMA/effector interaction are labelled as shown. **(B)** Schematic of HMA residues as for panel (A), but bound to AVR-PikE. **(C)** Superposition showing Pikp-HMA, Pikp-HMA^NK-KE^ and Pikm-HMA chains (coloured in blue, purple and yellow, respectively) bound to AVR-PikD. For clarity, only the Lys-261/262 side chain is shown. **(D)** Superposition as described before, but bound to AVR-PikE.

We found only limited structural perturbations at either of the other previously defined interfaces (interface 1 or 2 (22)) between the AVR-PikD or AVR-PikE effectors bound to Pikp-HMA or Pikp-HMA^NK-KE^ (**Figure 3 - figure supplement 2**). We therefore conclude that the effects of the Pikp^NK-KE^ mutant on protein function are mediated via altered interactions at interface 3.

### Mutation in AVR-Pik effectors at the engineered binding interface impacts in planta response and in vivo binding

To further confirm that the engineered binding interface is responsible for the expanded recognition of AVR-PikE and AVR-PikA by Pikp^NK-KE^, we used mutants in the effectors at interface 2 (AVR-PikD^H46E^) and interface 3 (AVR-PikD,E,A^E53R^), which have previously been shown to impact interactions and in planta responses in wild-type NLR alleles (22). We tested whether these mutants affected the cell death response in *N. benthamiana*, and interactions between effectors and Pikp-HMA^NK-KE^ (Y2H) and between effectors and full length Pikp^NK-KE^ (in planta immunoprecipitation).

Firstly, we investigated the impact of mutation at interface 2 using the AVR-PikD^H46E^ mutant. Similar to with Pikp, we found that cell death in *N. benthamiana* is essentially blocked when co-expressing Pikp^NK-KE^ with this mutant, suggesting that the engineered NLR is still reliant on this interface for response (**Figure 4A**). Intriguingly, Y2H shows that the AVR-PikD^H46E^ mutant displays some interaction with Pikp-HMA^NK-KE^ (**Figure 4B**), similar to this mutant’s interaction with Pikm-HMA (22), although this interaction is barely observed by co-immunoprecipitation with the full length NLR (**Figure 4C**).

**Figure 4.**
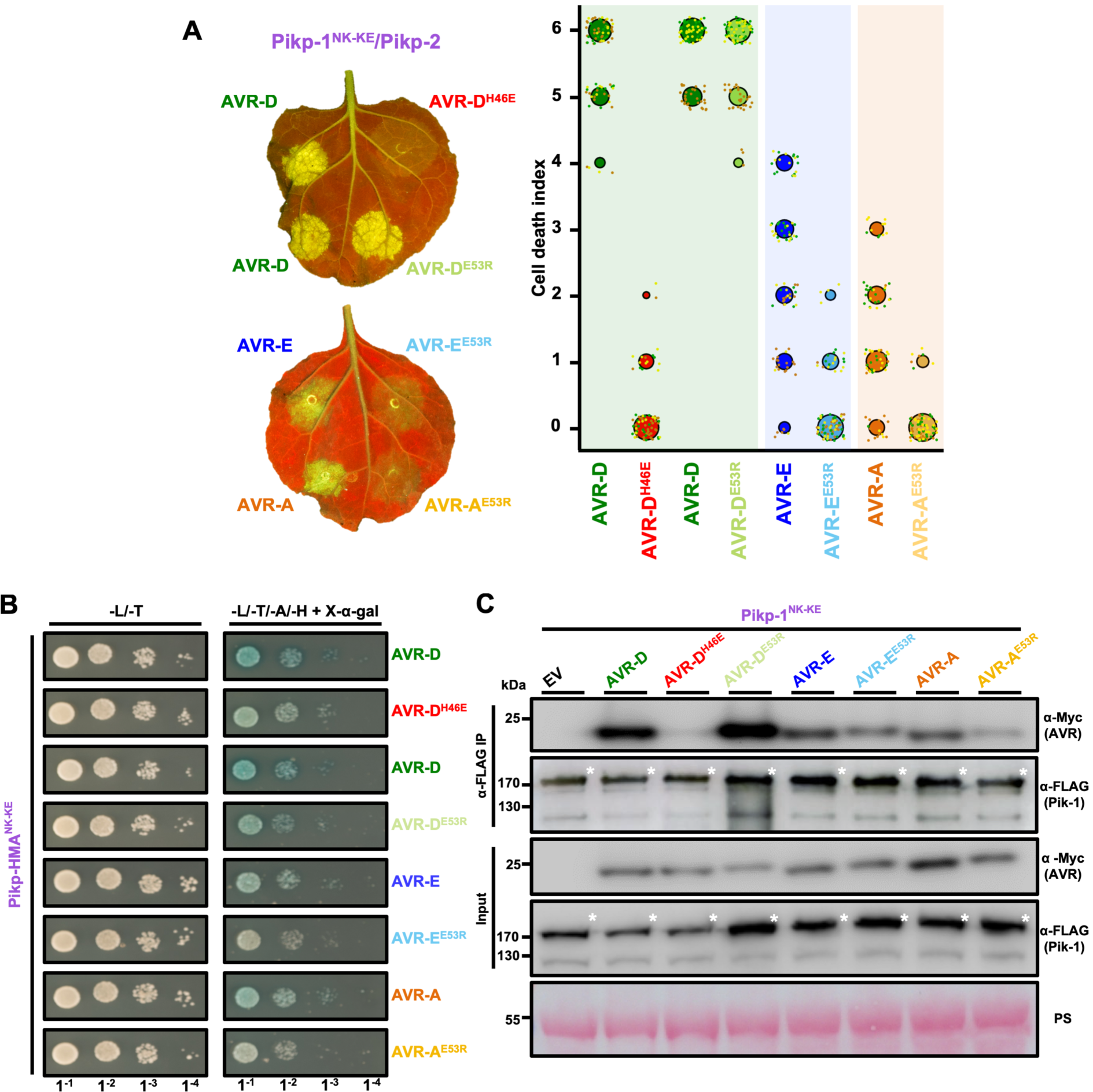
Mutation of AVR-Pik effectors at the engineered binding interface compromises binding and response. **(A)** Left - a representative leaf image showing Pikp-1^NK-KE^-mediated cell death to AVR-Pik variants and mutants as autofluorescence under UV light. Autofluorescence intensity is scored as in Figure 1. Right - Pikp^NK-KE^ cell death assay quantification is represented as dot plots. For each sample the data points are represented as dots with a distinct colour for each of the three biological replicates; these dots are jittered about the cell death score for visualisation purposes. The size of the central dot at each cell death value is proportional to the number of replicates of the sample with that score. The number of repeats was 90. **(B)** Yeast-Two-Hybrid assay of Pikp-HMA^NK-KE^ with AVR-Pik variants and mutants. Control plate for yeast growth is on the left, with the selective plate on the right for each combination of HMA/AVR-Pik. Growth and development of blue colouration in the selection plate are both indicative of protein:protein interaction. Each experiment was repeated a minimum of three times, with similar results. **(C)** Co-immunoprecipitation of full length Pikp-1^NK-KE^ with AVR-Pik variants and mutants. N-terminally 4xMyc tagged AVR-Pik effectors were transiently co-expressed with Pikp-1^NK-KE^:6xHis3xFLAG in *N. benthamiana* leaves. Immunoprecipitates (IPs) obtained with anti-FLAG antiserum, and total protein extracts, were probed with appropriate antisera. Each experiment was repeated at least three times, with similar results. The asterisks mark the Pik-1 band, PS = Ponceau Stain.

Secondly, we investigated the impact of mutations at interface 3 using the Glu53Arg (E53R) mutant in AVR-PikD, AVR-PikE and AVR-PikA. We found that the AVR-PikD^E53R^ mutant has essentially no effect on recognition of the effector by Pikp^NK-KE^ in *N. benthamiana*, and no effect on interaction with Pikp-HMA^NK-KE^, or full-length Pikp^NK-KE^ (**Figure 4A,B,C**). By contrast, the equivalent mutation in AVR-PikE and AVR-PikA essentially blocked the cell death response in *N. benthamiana*, reduced the binding to Pikp-HMA^NK-KE^ in Y2H (as shown by the reduced blue colouration) and a less intense band is observed for the effector following Pikp^NK-KE^ immunoprecipitation (**Figure 4A,B,C**). Expression of all proteins in yeast was confirmed by western blot (**Figure 4 – figure supplement 1**).

These results support that whilst interactions across interface 2 remain important for the Pikp^NK-KE^ interaction with AVR-Pik effectors, it is the altered interaction at interface 3, as observed in the structures, that is responsible for the expanded recognition profile of this engineered mutant.

## Discussion

Plants, including food crops, are under continuous threat from pathogens and pests, and new solutions to control disease are required. While largely elusive to date, engineering plant intracellular immune receptors (NLRs) has potential as a mechanism for improving disease resistance breeding (4, 6). NLR integrated domains are a particularly attractive target for protein engineering as they directly interact with pathogen effectors (or host effector targets). Further, where tested, binding affinities in vitro correlate with in planta immunity phenotypes (20, 22, 27), allowing biochemical and structural techniques to directly inform NLR design.

Here we show that the recognition profile of the rice NLR Pikp can be expanded to different AVR-Pik variants by engineering the binding interface between these proteins. This strengthens the hypothesis that tighter binding affinity between effectors and integrated HMA domains correlates with increased immune signalling in plants. This was previously shown for both natural alleles of Pik (20, 22), and also for Pia (27), but is now also shown for an engineered NLR. We propose this may be a general model for integrated domains that directly bind effectors.

Natural variation in Pik NLRs has given rise to different effector recognition profiles, and contribution from different binding interfaces was suggested to underpin this phenotype (22). In particular, a more favourable interaction at one interface (interface 3) in Pikm, compared to Pikp, was concluded to have evolved to compensate for changes in binding at a different site (interface 2). Here, through mutation of residues in Pikp (forming Pikp^NK-KE^), we have combined favourable interfaces from Pikp and Pikm into a single protein. This has resulted in an expanded recognition phenotype that also out-performs either of Pikp or Pikm effector binding and response in planta. While the Pikp^NK-KE^ mutant did not deliver a cell death response in *N. benthamiana* to the stealthy AVR-PikC effector variant, it did show a robust gain-of-binding in Y2H and in vitro, and showed very weak binding by in planta co-immunoprecipitation. We hypothesise that this gain-of-binding is as-yet not of sufficient strength, especially in the context of the full-length NLR, to trigger immune signalling. However, this work sets the scene for future interface engineering experiments that may further improve the response profiles of Pik NLRs to currently unrecognised effector variants. It also requires future work to test the disease resistance profile of *M. oryzae* strains carrying the different effector variants in rice expressing the engineered receptor.

The integrated HMA domain in the NLR RGA5 (the sensor of the Pia NLR pair in rice), binds to *M. oryzae* effectors AVR1-CO39 and AVR-Pia *via* a different interface, and it has been suggested that these binding sites are mutually exclusive (27). This raises the possibility that an HMA domain could be engineered to bind and respond to multiple effectors (27). Recently, the Pikp-HMA domain was shown to interact with AVR-Pia at the same interface as used by the RGA5-HMA domain, and this likely underpins partial resistance to *M. oryzae* expressing AVR-Pia in planta (31). This presents a starting point for using Pikp as a chassis for such studies. While it remains to be seen whether any such resurfaced HMA domain can bind to multiple effectors, these studies suggest this has potential as a novel approach.

Plant breeding is required to provide new genetic solutions to disease resistance in crops. This is necessary to limit the environmental and social damage caused by pesticides, and to deal with changes in climate and globalisation of agriculture that result in the spread of pathogens and pests into new environments (32–34). Classical breeding for disease resistance has been limited by issues such as linkage drag and hybrid incompatibility, as also seen in model plant species (35). Novel molecular approaches such as engineering “decoys” (12) and protein resurfacing, as described here, combined with modern transformation (36) and breeding pipelines (37), offers the opportunity for more targeted approaches to disease resistance breeding. These will complement other emerging technologies in NLR identification (38) and NLR stacking (4) as methods to develop improved crops for the future.

## Accession codes

Protein structures, and the data used to derive these, have been deposited at the Protein DataBank (PDB) with accession codes 6R8K (Pikp-HMA^NK-KE^/AVR-PikD) and 6R8M (Pikp-HMA^NK-KE^/AVR-PikE).

## Acknowledgements

This work was supported by the BBSRC (grants BB/J004553, BB/P012574, BB/M02198X), the ERC (proposal 743165), the John Innes Foundation, the Gatsby Charitable Foundation, and JSPS KAKENHI 15H05779. We thank the Diamond Light Source, UK (beamline i03 under proposal MX13467) for access to X-ray data collection facilities. We also thank David Lawson and Clare Stevenson (JIC X-ray Crystallography/Biophysical Analysis Platform) for help with protein structure determination and SPR.

## Materials and Methods

### Gene cloning

For in vitro studies, Pikp-HMA^NK-KE^ (encompassing residues 186 to 263) was amplified from WT Pikp-HMA by introducing the mutations in the reverse primer, followed by cloning into pOPINM (39). Wild-type Pikp-HMA, Pikm-HMA, and AVR-Pik expression constructs used in this study are as described in (22).

For Y2H, we cloned Pikp-HMA^NK-KE^ (as above) into pGBKT7 using an In-Fusion cloning kit (Takara Bio USA), following the manufacture’s protocol. Wild-type Pikp-HMA domain in pGBKT7 and AVR-Pik effector variants in pGADT7 used were generated as described in (22).

For protein expression in planta, Pikp-HMA^NK-KE^ domain was generated using site directed mutagenesis by introducing the mutations in the reverse primer. This domain was then assembled into a full-length construct using Golden Gate cloning (40) and into the plasmid pICH47742 with a C-terminal 6xHis/3xFLAG tag. Expression was driven by the *A. tumefaciens* Mas promoter and terminator. Full-length Pikp-1, Pikp-2, and AVR-Pik variants used were generated as described in (22).

All DNA constructs were verified by sequencing.

### Expression and purification of proteins for in vitro binding studies

pOPINM constructs encoding Pikp-HMA, Pikm-HMA and Pikp-HMA^NK-KE^ were produced in *E. coli* SHuffle cells (41) using the same protocol described in (22). Cell cultures were grown in auto induction media (42) at 30°C for 5 – 7hrs and then at 16°C overnight. Cells were harvested by centrifugation and re-suspended in 50 mM Tris-HCl pH7.5, 500 mM NaCl, 50 mM Glycine, 5% (vol/vol) glycerol, 20 mM imidazole supplemented with EDTA-free protease inhibitor tablets (Roche). Cells were sonicated and, following centrifugation at 40000xg for 30 min, the clarified lysate was applied to a Ni^2+^-NTA column connected to an AKTA Xpress purification system (GE Healthcare). Proteins were step-eluted with elution buffer (50 mM Tris-HCl pH7.5, 500 mM NaCl, 50 mM Glycine, 5% (vol/vol) glycerol, 500 mM imidazole) and directly injected onto a Superdex 75 26/60 gel filtration column pre-equilibrated 20mM HEPES pH 7.5, 150 mM NaCl. Purification tags were removed by incubation with 3C protease (10 μg/mg fusion protein) followed by passing through tandem Ni^2+^-NTA and MBP Trap HP columns (GE Healthcare). The flow-through was concentrated as appropriate and loaded on a Superdex 75 26/60 gel filtration column for final purification and buffer exchange into 20 mM HEPES pH 7.5, 150 mM NaCl.

AVR-Pik effectors, with either a 3C protease-cleavable N-terminal SUMO or MBP tag, and a non-cleavable C-terminal 6xHis tag, were produced in and purified from *E. coli* SHuffle cells as previously described (20, 22). All protein concentrations were determined using a Direct Detect^®^ Infrared Spectrometer (Merck).

### Co-expression and purification of Pik-HMA/AVR-Pik effectors for crystallisation

Pikp-HMA^NK-KE^ was co-expressed with AVR-PikD or AVR-PikE effectors in *E. coli* SHuffle cells following co-transformation of pOPINM:Pikp-HMA^NK-KE^ and pOPINA:AVR-PikD/E (which were prepared as described in (22)). Cells were grown in autoinduction media (supplemented with both carbenicillin and kanamycin), harvested, and processed as described in (22). Protein concentrations were measured using a Direct Detect® Infrared Spectrometer (Merck).

### Protein-protein interaction: Yeast-2-hybrid analyses

To detect protein–protein interactions between Pik-HMAs and AVR-Pik effectors by Yeast Two-Hybrid, we used the Matchmaker® Gold System (Takara Bio USA). Plasmid DNA encoding Pikp-HMA^NK-KE^ in pGBKT7, generated in this study, was co-transformed into chemically competent Y2HGold cells (Takara Bio, USA) with the individual AVR-Pik variants or mutants in pGADT7 described previously (22). Single colonies grown on selection plates were inoculated in 5 ml of SD^-Leu-Trp^ overnight at 30°C. Saturated culture was then used to make serial dilutions of OD_600_ 1, 1^-1^, 1^-2^, 1^-3^, respectively. 5 μl of each dilution was then spotted on a SD^-Leu-Trp^ plate as a growth control, and on a SD^-Leu-Trp-Ade-His^ plate containing X-α-gal. Plates were imaged after incubation for 60 - 72 hr at 30°C. Each experiment was repeated a minimum of 3 times, with similar results.

To confirm protein expression in yeast, total protein extracts from transformed colonies were produced by boiling the cells 10 minutes in LDS Runblue® sample buffer. Samples were centrifugated and the supernatant was subjected to SDS-PAGE gels prior to western blotting. The membranes were probed with anti-GAL4 DNA-BD (Sigma) for the HMA domains in pGBKT7 and anti-GAL4 activation domain (Sigma) antibodies for the AVR-Pik effectors in pGADT7.

### Protein-protein interaction: Surface plasmon resonance

Surface plasmon resonance (SPR) experiments were performed on a Biacore T200 system (GE Healthcare) using an NTA sensor chip (GE Healthcare). The system was maintained at 25°C, and a flow rate of 30 μl/min was used. All proteins were prepared in SPR running buffer (20 mM HEPES pH 7.5, 860 mM NaCl, 0.1% Tween 20). C-terminally 6xHis-tag AVR-Pik variants were immobilised on the chip, giving a response of 200 ± 100. The sensor chip was regenerated between each cycle with an injection of 30 μl of 350 mM EDTA.

For all the assays, the level of binding was expressed as a percentage of the theoretical maximum response (R_max_) normalized for the amount of ligand immobilized on the chip. The cycling conditions were the same as used in (22). For each measurement, in addition to subtracting the response in the reference cell, a further buffer-only subtraction was made to correct for bulk refractive index changes or machine effects (43). SPR data was exported and plotted using R v3.4.3 (https://www.r-project.org/) and the function ggplot2 (Wickham, H., 2009). Each experiment was repeated a minimum of 3 times, including internal repeats, with similar results.

### Protein-protein interaction: In planta co-immunoprecipitation (Co-IP)

Transient gene-expression in planta for Co-IP was performed by delivering T-DNA constructs with *Agrobacterium tumefaciens* GV3101 strain into 4-week old *N. benthamiana* plants grown at 22–25°C with high light intensity. *A. tumefaciens* strains carrying Pikp-1 or Pikp -1^NK-KE^ were mixed with strains carrying the AVR-Pik effector, at OD_600_ 0.2 each, in agroinfiltration medium (10 mM MgCl_2_, 10 mM 2-(N-morpholine)-ethanesulfonic acid (MES), pH5.6), supplemented with 150 μM acetosyringone. For detection of complexes in planta, leaf tissue was collected 3 days post infiltration (dpi), frozen, and ground to fine powder in liquid nitrogen using a pestle and mortar. Leaf powder was mixed with 2 times weight/volume ice-cold extraction buffer (10% glycerol, 25 mM Tris pH 7.5, 1 mM EDTA, 150 mM NaCl, 2% w/v PVPP, 10 mM DTT, 1x protease inhibitor cocktail (Sigma), 0.1% Tween 20 (Sigma)), centrifuged at 4,200g/4°C for 20-30 min, and the supernatant was passed through a 0.45μm Minisart® syringe filter. The presence of each protein in the input was determined by SDS-PAGE/western blot. Pik-1 and AVR-Pik effectors were detected probing the membrane with anti-FLAG M2 antibody (SIGMA) and anti c-Myc monoclonal antibody (Santa Cruz), respectively. For immunoprecipitation, 1.5ml of filtered plant extract was incubated with 30 μl of M2 anti-FLAG resin (Sigma) in a rotatory mixer at 4°C. After three hours, the resin was pelleted (800g, 1 min) and the supernatant removed. The pellet was washed and resuspended in 1ml of IP buffer (10% glycerol, 25 mM Tris pH 7.5, 1 mM EDTA, 150 mM NaCl, 0.1% Tween 20 (Sigma)) and pelleted again by centrifugation as before. Washing steps were repeated 5 times. Finally, 30 μl of LDS Runblue® sample buffer was added to the agarose and incubated for 10 min at 70°C. The resin was pelleted again, and the supernatant loaded on SDS-PAGE gels prior to western blotting. Membranes were probed with anti-FLAG M2 (Sigma) and anti c-Myc (Santa Cruz) monoclonal antibodies.

#### *N. benthamiana* cell death assays

*A. tumefaciens* GV3101 cells, transformed with the relevant constructs, were spot inoculated on 4-weeks old *N. benthamiana* leaves using a needleless syringe. Strains carrying Pikp-1 or Pikp-1^NK-KE^ were mixed with Pikp-2, AVR-Pik effectors and P19 at OD_600_ 0.4, 0.4, 0.6 and 0.1, respectively. Detached leaves were imaged at 5 dpi from the abaxial side of the leaves for UV images. Images are representative of three independent experiments, with internal repeats. The cell death index used for scoring is as presented previously (20). Dotplots were generated using R v3.4.3 (https://www.r-project.org/) and the graphic package ggplot2 (Wickham, H., 2009). The size of the centre dot at each cell death value is directly proportional to the number of replicates in the sample with that score. All individual data points are represented as dots.

### Crystallization, data collection and structure solution

For crystallization, Pikp-HMA^NK-KE^ in complex with AVR-PikD or AVR-PikE were concentrated following gel filtration. Sitting drop vapor diffusion crystallization trials were set up in 96 well plates, using an Oryx nano robot (Douglas Instruments, United Kingdom). Plates were incubated at 20°C, and crystals typically appeared after 24 - 48 hours. For data collection, all crystals were harvested from the Morpheus® HT-96 screen (Molecular Dimensions), and snap-frozen in liquid nitrogen. Crystals used for data collection appeared from the following conditions: (i) Pikp-HMA^NK-KE^/AVR-PikD (10 mg/ml), Morpheus® HT-96 condition D4 [0.12 M Alcohols (0.2 M 1,6-Hexanediol; 0.2 M 1-Butanol; 0.2 M 1,2-Propanediol; 0.2 M 2-Propanol; 0.2 M 1,4-Butanediol; 0.2 M 1,3-Propanediol); 0.1 M Buffer system 1 (1 M Imidazole; MES monohydrate (acid)) pH 6.5; 50% v/v Precipitant mix 4 (25%v/v MPD; 25%v/v PEG 1000; 25%v/v PEG3350)]; (ii) Pikp-HMA^NK-KE^/AVR-PikE (15 mg/ml), Morpheus® HT-96 condition A8 [0.06M Divalents (0.3 M Magnesium chloride hexahydrate; 0.3 M Calcium chloride dihydrate); 0.1M Buffer system 2 (Sodium HEPES; MOPS (acid)) pH 7.5; 37.5%v/v Precipitant mix 4 (25%v/v MPD; 25%v/v PEG 1000; 25%v/v PEG3350)].

X-ray data sets were collected at the Diamond Light Source using beamline i03 (Oxford, UK). The data were processed using the xia2 pipeline (44) and CCP4 (45). The structures were solved by molecular replacement using PHASER (46) and the coordinates of AVR-PikD and a monomer of Pikp-HMA from PDB entry 6G10. The final structures were obtained through iterative cycles of manual rebuilding and refinement using COOT (47) and REFMAC5 (48), as implemented in CCP4 (45). Structures were validated using the tools provided in COOT and MOLPROBITY (49).

---

**Figure 1 figure supplement 1.**
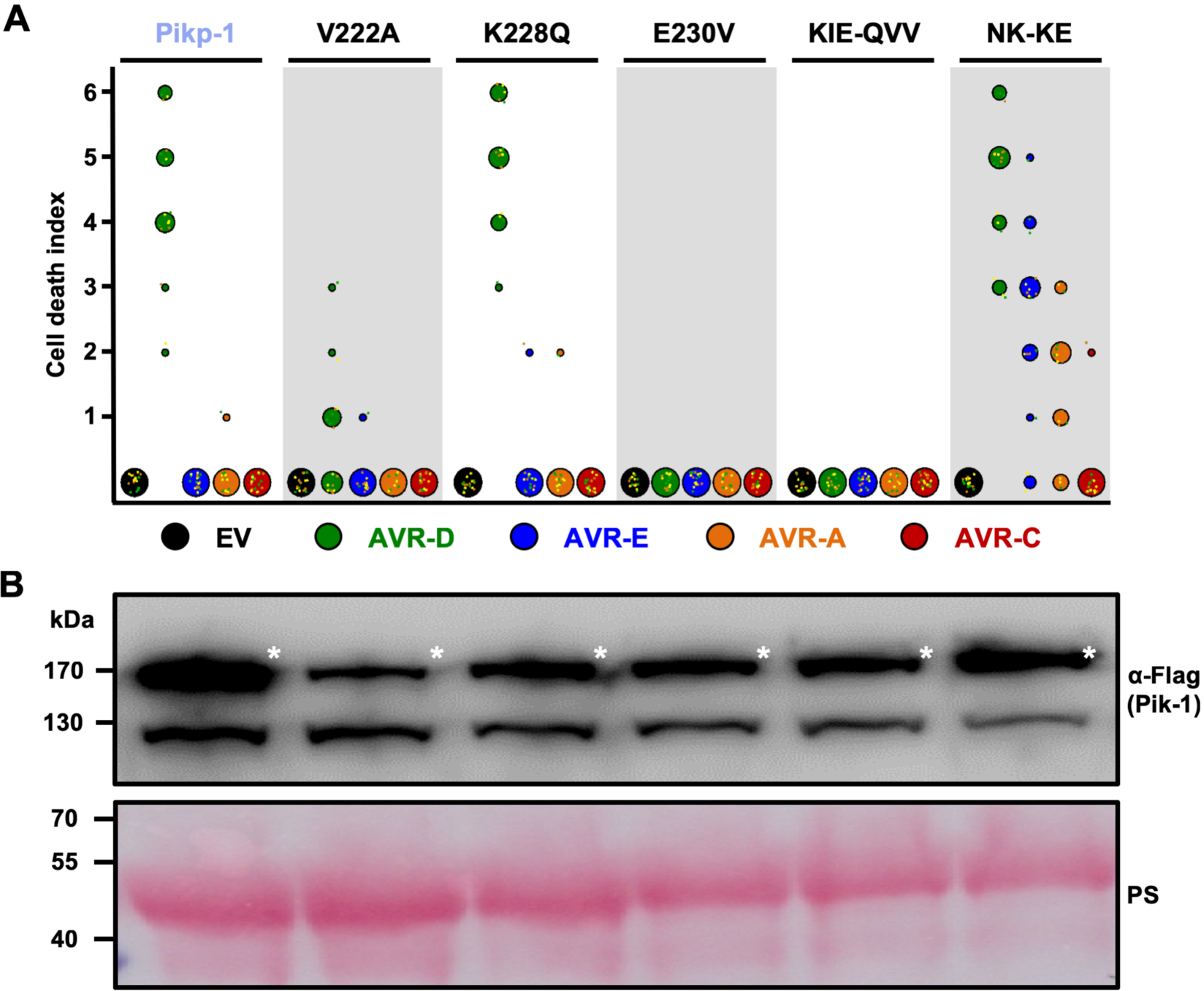
Mutations at interface 2 of Pikp-1 HMA domain compromise response to AVR-Pik effectors. **(A)** Cell death assay scoring represented as dot plots for Pikp-1 mutants on HMA interface 2 and 3. For each sample, all the data points are represented as dots with a distinct colour for each of the three biological replicates; these dots are jittered about the cell death score for visualisation purposes. The size of the central dot at each cell death value is proportional to the number of replicates of the sample with that score. The number of repeats was 18 for each mutant. **(B)** Western blot analysis confirming similar levels of Pik-1 protein accumulation in *N. benthamiana*. The asterisks mark the Pik-1 band, PS = Ponceau Stain.

**Figure 1 figure supplement 2.**
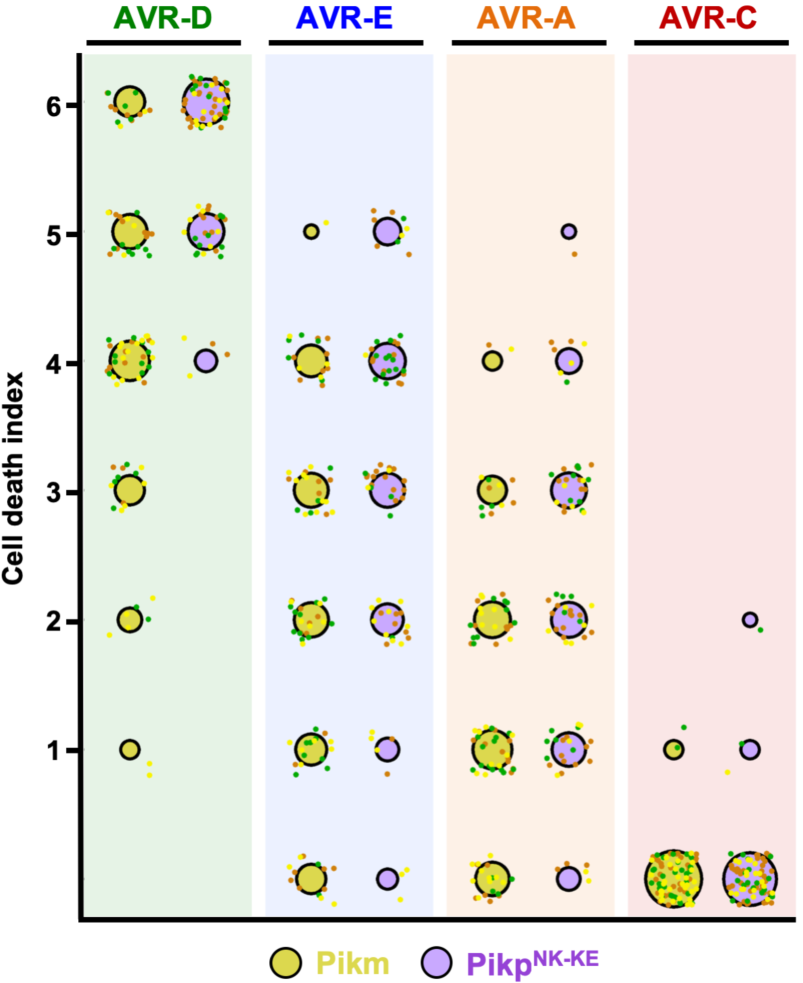
Pikp^NK-KE^ shows a qualitatively stronger response to AVR-Pik effectors compared to Pikm. Cell death assay scoring represented as dot plots for Pikm and Pikp^NK-KE^ (yellow and purple respectively). The number of repeats was 80 and 90 for Pikp^NK-KE^ and Pikm, respectively. For each sample, all the data points are represented as dots with a distinct colour for each of the three biological replicates; these dots are jittered about the cell death score for visualisation purposes. The size of the central dot at each cell death value is proportional to the number of replicates of the sample with that score. Data for Pikm has been previously shown (22), but was acquired at the same time as Pikp^NK-KE^.

**Figure 2 figure supplement 1.**
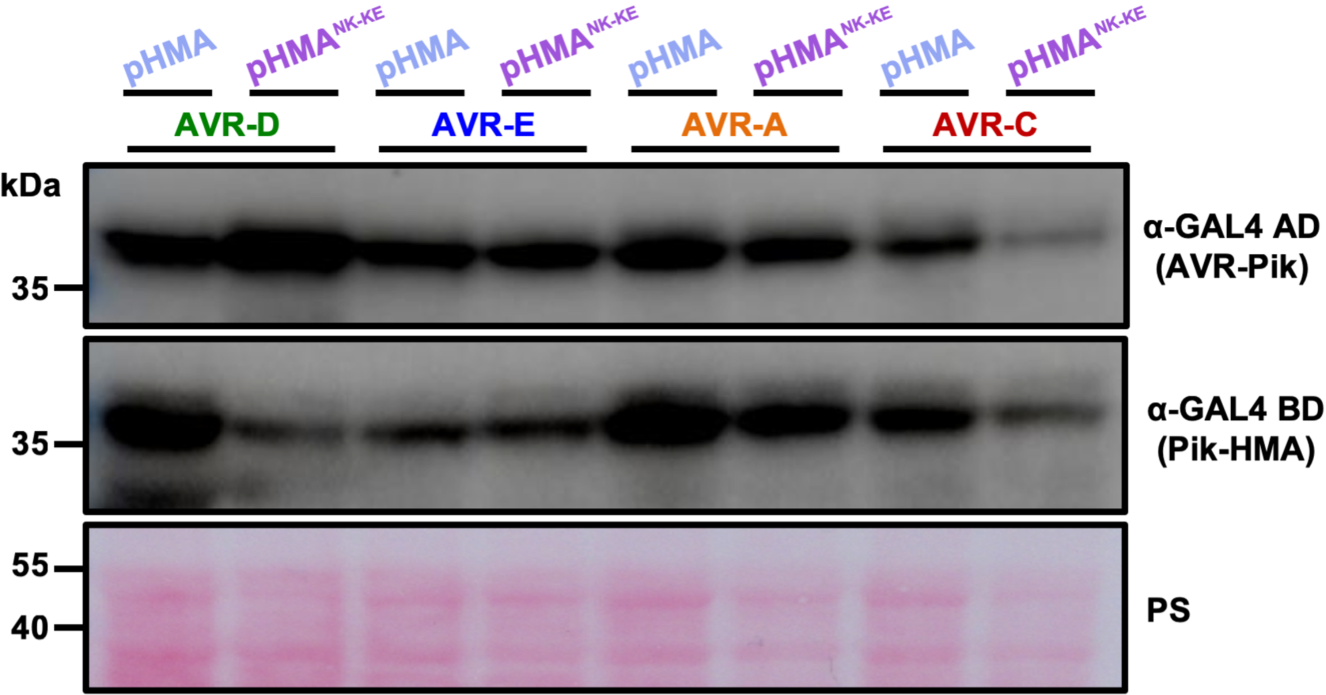
Western blot confirming accumulation of proteins in yeast. Yeast lysate was probed for the expression of HMA domains with anti-GAL4 DNA binding domain (BD) and AVR-Pik effectors anti-GAL4 activation domain (AD). Total extract was coloured with Ponceau Stain (PS). The experiment was repeated a minimum of 3 times, with similar results. PS = Ponceau Stain.

**Figure 2 figure supplement 2.**
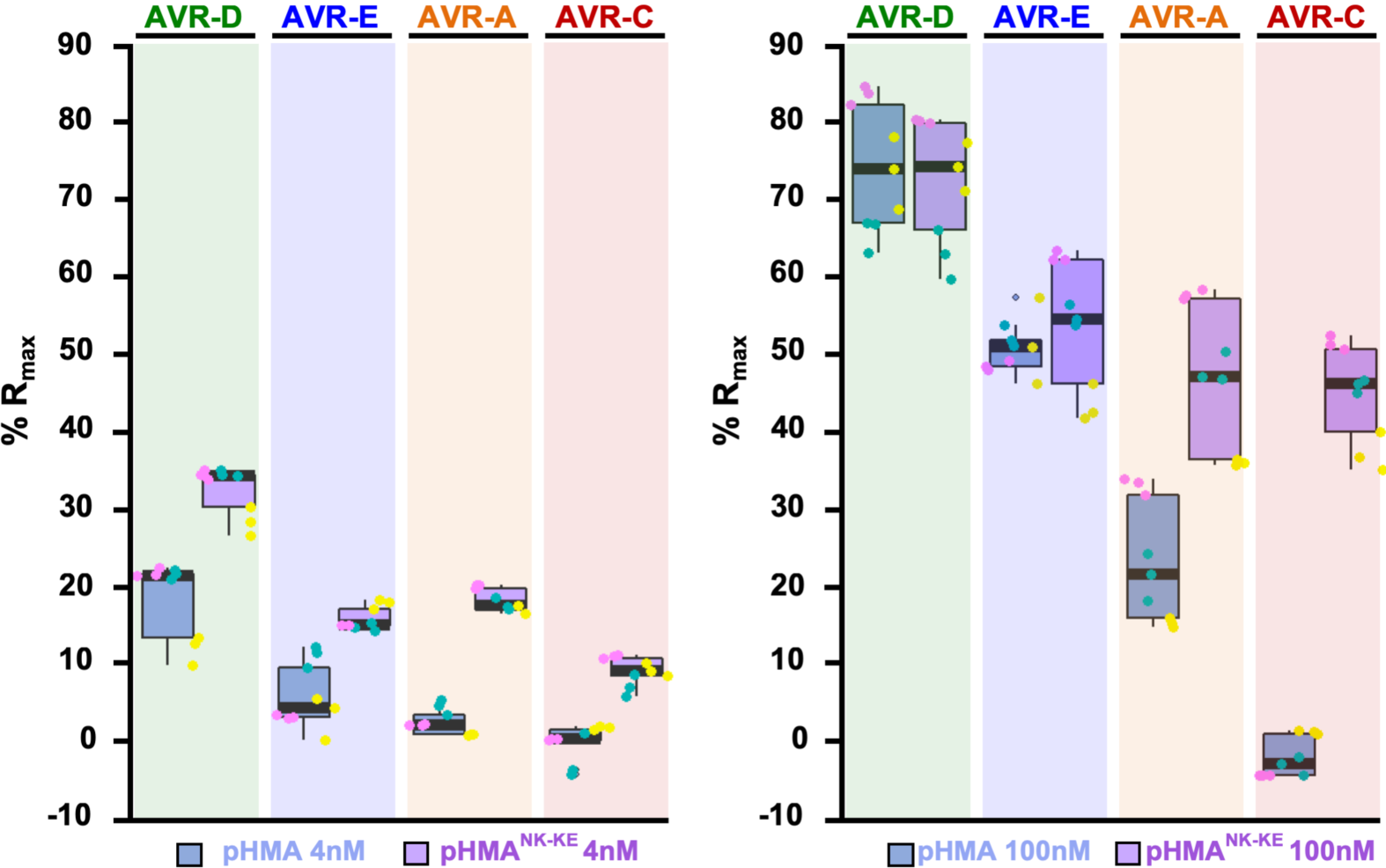
Binding of the Pikp-HMA^NK-KE^ domain to the AVR-Pik effectors is consistently higher compared to Pikp-HMA. %R_max_ of Pikp-HMA and Pikp-HMA^NK-KE^ with the AVR-Pik effectors alleles with HMA concentrations of 4nM (left) and 100nM (right).

**Figure 2 figure supplement 3.**
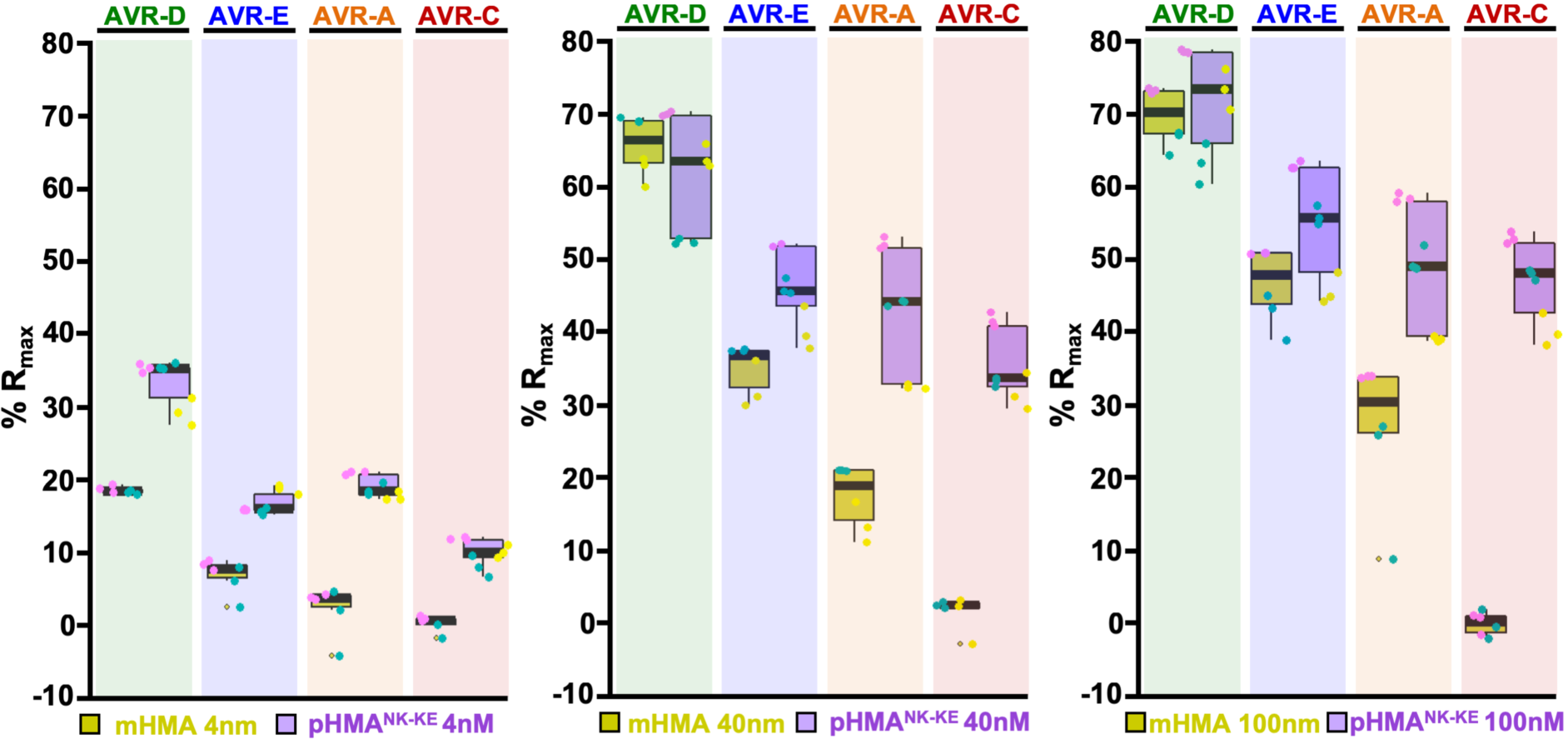
Binding of the Pikp-HMA^NK-KE^ domain to the AVR-Pik effectors is consistently higher compared to Pikm-HMA. Surface plasmon resonance %R_max_ values for Pikm-HMA and Pikp-HMA^NK-KE^ with the AVR-Pik effectors alleles. Pikm-HMA and Pikp-HMA^NK-KE^ results are represented by yellow and purple boxes, respectively. The centre line represents the median, the box limits are the upper and lower quartiles, the whiskers are the 1.5× interquartile range and all the data points are represented as dots with distinct colours for each biological replicate. For each experiment, we performed at least two biological replicates with three internal repeats. Results using HMA protein concentration of 4, 40 and 100 nM are plotted in the left, middle and right panels, respectively.

**Figure 3 figure supplement 1.**
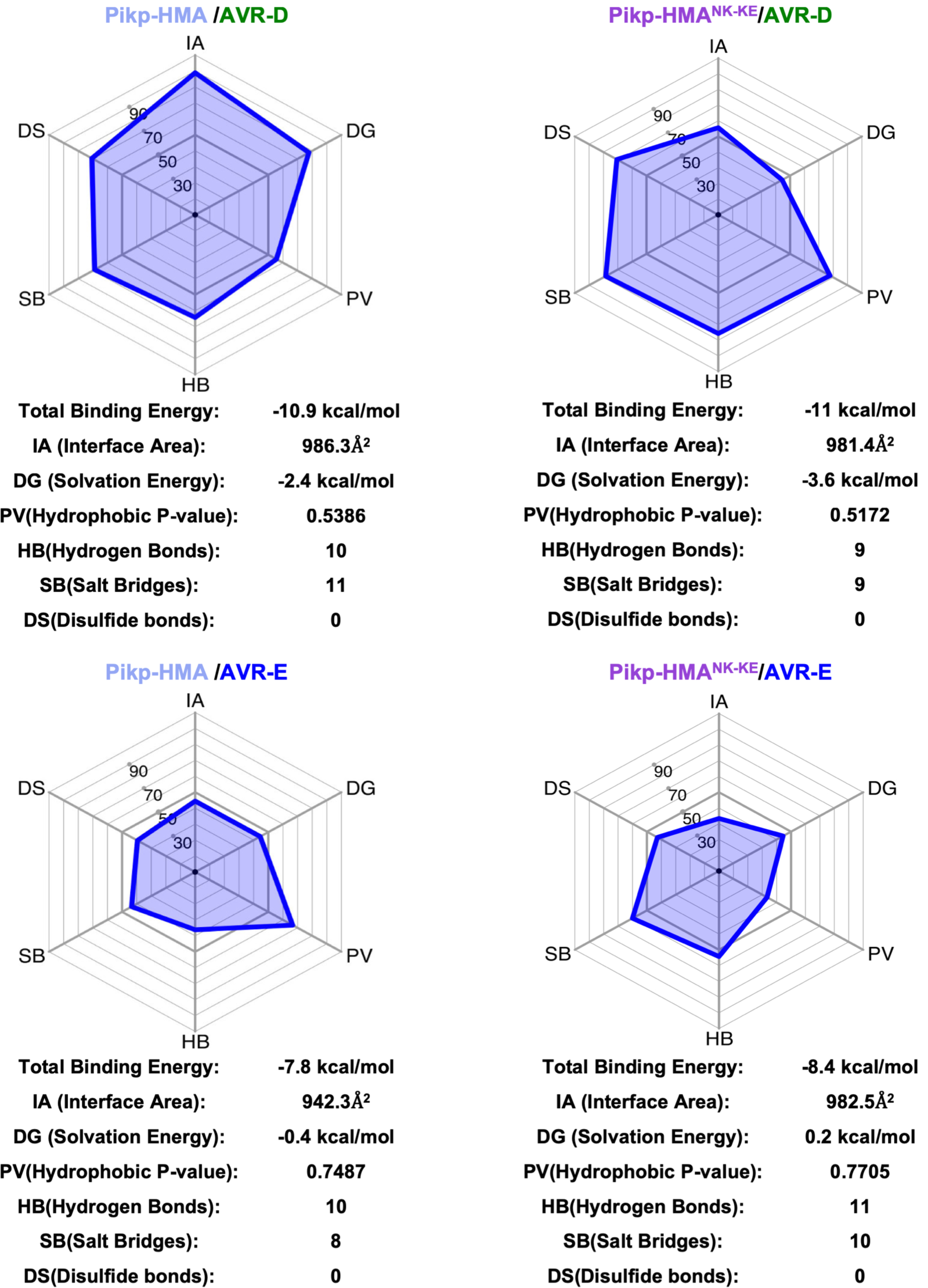
Analyses of Pikp-HMA and Pikp-HMA^NK-KE^ interfaces with AVR-PikD or AVR-PikE using QtPISA. For each complex (as labelled), the QtPISA interaction radar was generated with reference parameter “Total Binding Energy”. The internal area of the polygon represents the Total Binding Energy. The values obtained for each key interface parameter are shown on the axes, with these scores based on statistical analysis of all interfaces found in the Protein DataBank. The abbreviations in the radars are defined in the tables underneath each panel.

**Figure 3 figure supplement 2.**
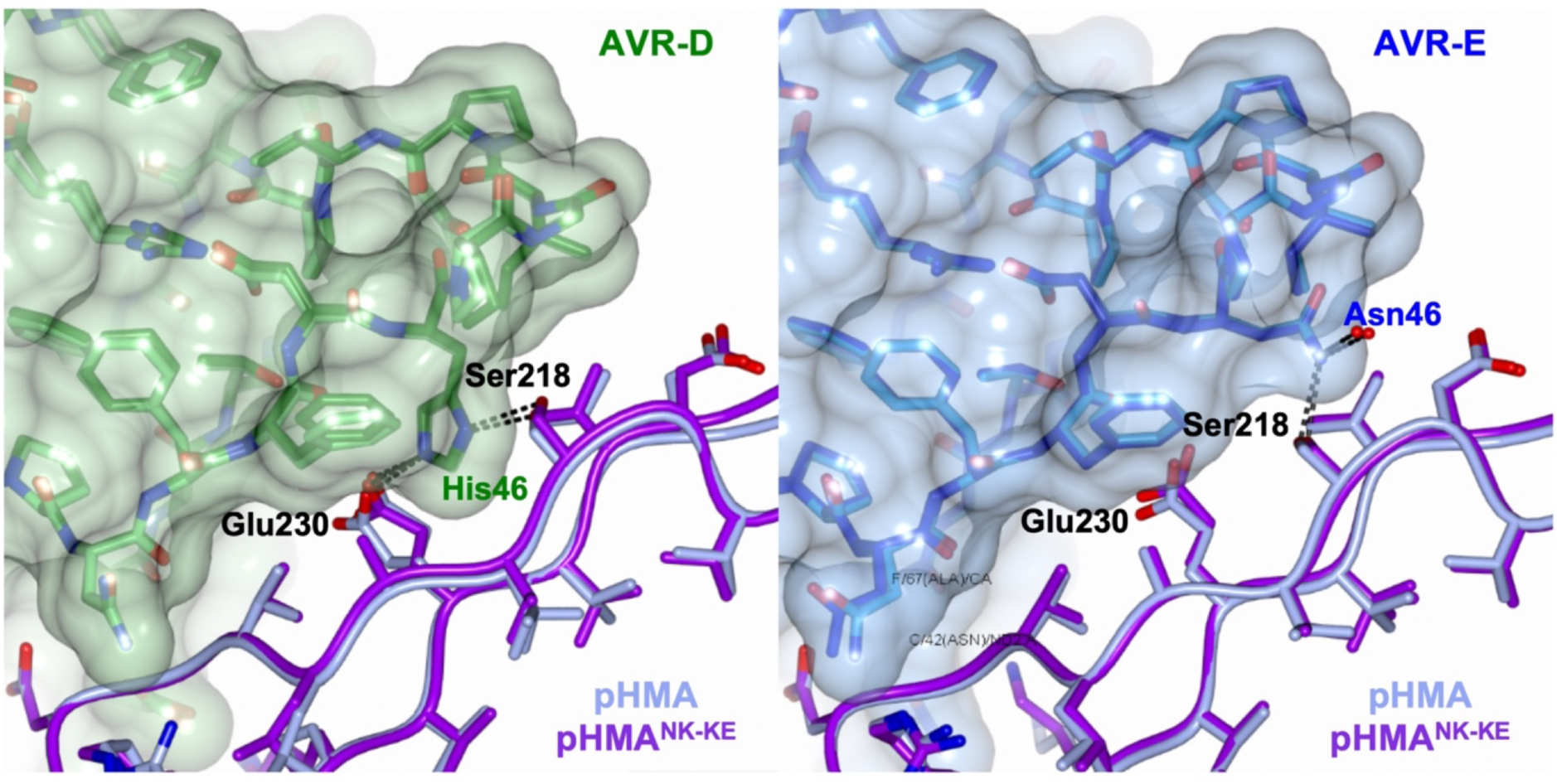
Interface 2 is essentially identical in the complexes of Pikp-HMA and Pikp-HMA^NK-KE^ bound to AVR-PikD or AVR-PikE. Schematic view of the conformations adopted by Pikp-HMA and Pikp-HMA^NK-KE^ at interface 2 in complex with AVR-PikD or AVR-PikE. In each panel, the effector is shown as sticks with the molecular surface also shown and coloured as labelled. Pik-HMA residues are coloured as labelled and shown in the Cα-worm with side-chain representation. The structures were overlaid on the effectors. **(A)** Pikp-HMA and Pikp-HMA^NK-KE^ (coloured in blue and purple, respectively) bound to AVR-PikD (light and dark green). **(B)** Pikp-HMA and Pikp-HMA^NK-KE^bound to AVR-PikE (light and dark blue).

**Figure 4 figure supplement 1.**
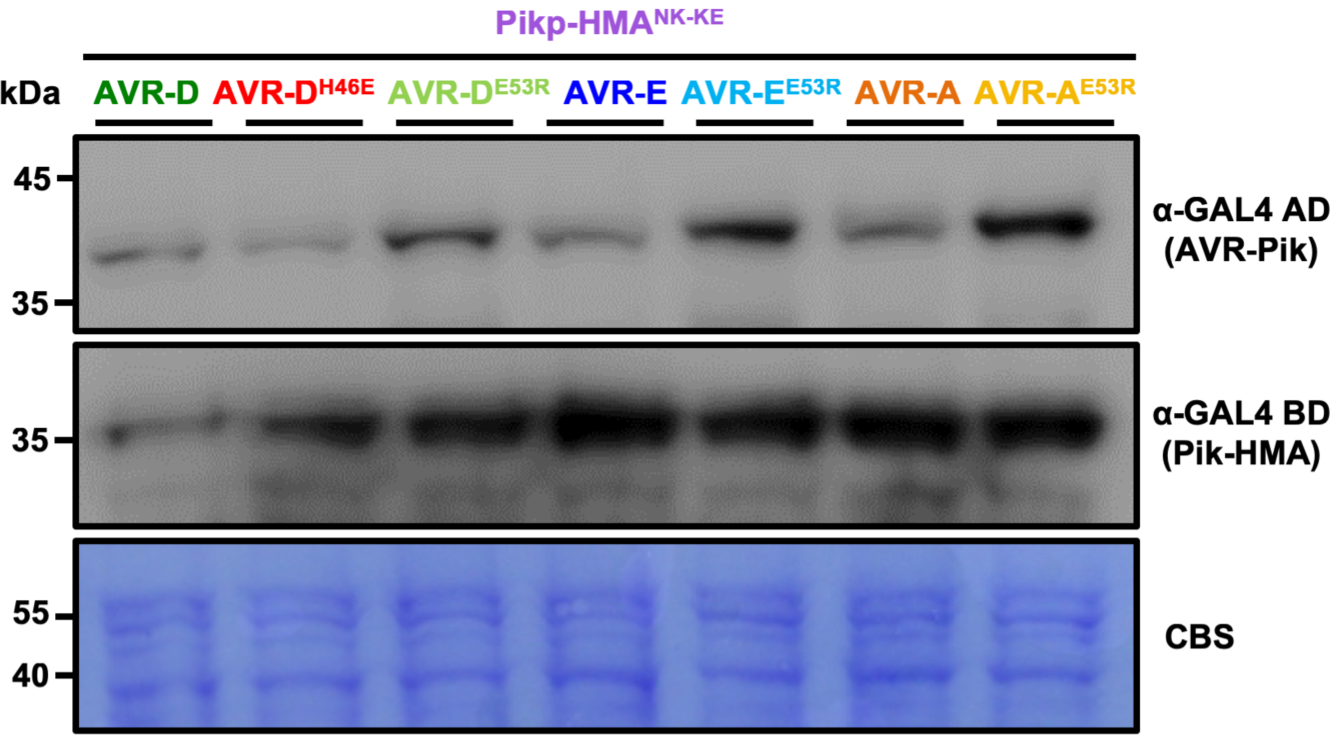
Western blot analysis confirming accumulation of proteins in yeast. Yeast lysate was probed for the expression of HMA domain with anti-GAL4 DNA binding domain (BD) and AVR-Pik effectors anti-GAL4 activation domain (AD). The experiment was repeated a minimum of 3 times, with similar results. CBS = Coomassie Blue Stain.

## References

1. Chapman, A. M., and McNaughton, B. R. (2016) Scratching the Surface: Resurfacing Proteins to Endow New Properties and Function. Cell Chem Biol 23, 543–553

2. Kourelis, J., and van der Hoorn, R. A. L. (2018) Defended to the Nines: 25 Years of Resistance Gene Cloning Identifies Nine Mechanisms for R Protein Function. Plant Cell 30, 285–299

3. Cesari, S. (2018) Multiple strategies for pathogen perception by plant immune receptors. New Phytol 219, 17–24

4. Dangl, J. L., Horvath, D. M., and Staskawicz, B. J. (2013) Pivoting the plant immune system from dissection to deployment. Science 341, 746–751

5. Yoshida, K., Saunders, D. G., Mitsuoka, C., Natsume, S., Kosugi, S., Saitoh, H., Inoue, Y., Chuma, I., Tosa, Y., Cano, L. M., Kamoun, S., and Terauchi, R. (2016) Host specialization of the blast fungus Magnaporthe oryzae is associated with dynamic gain and loss of genes linked to transposable elements. BMC Genomics 17, 370

6. Rodriguez-Moreno, L., Song, Y., and Thomma, B. P. (2017) Transfer and engineering of immune receptors to improve recognition capacities in crops. Curr Opin Plant Biol 38, 42–49

7. Segretin, M. E., Pais, M., Franceschetti, M., Chaparro-Garcia, A., Bos, J. I., Banfield, M. J., and Kamoun, S. (2014) Single amino acid mutations in the potato immune receptor R3a expand response to Phytophthora effectors. Mol Plant Microbe Interact

8. Giannakopoulou, A., Steele, J. F., Segretin, M. E., Bozkurt, T. O., Zhou, J., Robatzek, S., Banfield, M. J., Pais, M., and Kamoun, S. (2015) Tomato I2 Immune Receptor Can Be Engineered to Confer Partial Resistance to the Oomycete Phytophthora infestans in Addition to the Fungus Fusarium oxysporum. Mol Plant Microbe Interact 28, 1316–1329

9. Harris, C. J., Slootweg, E. J., Goverse, A., and Baulcombe, D. C. (2013) Stepwise artificial evolution of a plant disease resistance gene. Proc Natl Acad Sci U S A 110, 21189–21194

10. Carter, M. E., Helm, M., Chapman, A., Wan, E., Restrepo Sierra, A. M., Innes, R., Bogdanove, A. J., and Wise, R. P. (2018) Convergent evolution of effector protease recognition by Arabidopsis and barley. Mol Plant Microbe Interact

11. Helm, M., Qi, M., Sarkar, S., Yu, H., Whitham, S. A., and Innes, R. W. (2019) Engineering a Decoy Substrate in Soybean to Enable Recognition of the Soybean Mosaic Virus NIa Protease. Mol Plant Microbe Interact

12. Kim, S. H., Qi, D., Ashfield, T., Helm, M., and Innes, R. W. (2016) Using decoys to expand the recognition specificity of a plant disease resistance protein. Science 351, 684–687

13. Takken, F. L., and Goverse, A. (2012) How to build a pathogen detector: structural basis of NB-LRR function. Curr Opin Plant Biol 15, 375–384

14. Kroj, T., Chanclud, E., Michel-Romiti, C., Grand, X., and Morel, J. B. (2016) Integration of decoy domains derived from protein targets of pathogen effectors into plant immune receptors is widespread. New Phytol 210, 618–626

15. Sarris, P. F., Cevik, V., Dagdas, G., Jones, J. D., and Krasileva, K. V. (2016) Comparative analysis of plant immune receptor architectures uncovers host proteins likely targeted by pathogens. BMC Biol 14, 8

16. Bailey, P. C., Schudoma, C., Jackson, W., Baggs, E., Dagdas, G., Haerty, W., Moscou, M., and Krasileva, K. V. (2018) Dominant integration locus drives continuous diversification of plant immune receptors with exogenous domain fusions. Genome Biol 19, 23

17. Fujisaki, K. A. A.; Kanzaki, E.; Ito, K.; Utsushi, H.; Saitoh, H.; Białas, A.; Banfield, M.J.; Kamoun, S.; Terauchi, R. (2017) An unconventional NOI/RIN4 domain of a rice NLR protein binds host EXO70 protein to confer fungal immunity. bioRxiv

18. Cesari, S., Bernoux, M., Moncuquet, P., Kroj, T., and Dodds, P. N. (2014) A novel conserved mechanism for plant NLR protein pairs: the “integrated decoy” hypothesis. Front Plant Sci 5, 606

19. Le Roux, C., Huet, G., Jauneau, A., Camborde, L., Tremousaygue, D., Kraut, A., Zhou, B., Levaillant, M., Adachi, H., Yoshioka, H., Raffaele, S., Berthome, R., Coute, Y., Parker, J. E., and Deslandes, L. (2015) A receptor pair with an integrated decoy converts pathogen disabling of transcription factors to immunity. Cell 161, 1074–1088

20. Maqbool, A., Saitoh, H., Franceschetti, M., Stevenson, C. E., Uemura, A., Kanzaki, H., Kamoun, S., Terauchi, R., and Banfield, M. J. (2015) Structural basis of pathogen recognition by an integrated HMA domain in a plant NLR immune receptor. Elife 4

21. Sarris, P. F., Duxbury, Z., Huh, S. U., Ma, Y., Segonzac, C., Sklenar, J., Derbyshire, P., Cevik, V., Rallapalli, G., Saucet, S. B., Wirthmueller, L., Menke, F. L. H., Sohn, K. H., and Jones, J. D. G. (2015) A Plant Immune Receptor Detects Pathogen Effectors that Target WRKY Transcription Factors. Cell 161, 1089–1100

22. De la Concepcion, J. C., Franceschetti, M., Maqbool, A., Saitoh, H., Terauchi, R., Kamoun, S., and Banfield, M. J. (2018) Polymorphic residues in rice NLRs expand binding and response to effectors of the blast pathogen. Nat Plants 4, 576–585

23. Bialas, A., Zess, E. K., De la Concepcion, J. C., Franceschetti, M., Pennington, H. G., Yoshida, K., Upson, J. L., Chanclud, E., Wu, C. H., Langner, T., Maqbool, A., Varden, F. A., Derevnina, L., Belhaj, K., Fujisaki, K., Saitoh, H., Terauchi, R., Banfield, M. J., and Kamoun, S. (2018) Lessons in Effector and NLR Biology of Plant-Microbe Systems. Mol Plant Microbe Interact 31, 34–45

24. Eitas, T. K., and Dangl, J. L. (2010) NB-LRR proteins: pairs, pieces, perception, partners, and pathways. Curr Opin Plant Biol 13, 472–477

25. Cesari, S., Thilliez, G., Ribot, C., Chalvon, V., Michel, C., Jauneau, A., Rivas, S., Alaux, L., Kanzaki, H., Okuyama, Y., Morel, J. B., Fournier, E., Tharreau, D., Terauchi, R., and Kroj, T. (2013) The rice resistance protein pair RGA4/RGA5 recognizes the Magnaporthe oryzae effectors AVR-Pia and AVR1-CO39 by direct binding. Plant Cell 25, 1463–1481

26. Ortiz, D., de Guillen, K., Cesari, S., Chalvon, V., Gracy, J., Padilla, A., and Kroj, T. (2017) Recognition of the Magnaporthe oryzae Effector AVR-Pia by the Decoy Domain of the Rice NLR Immune Receptor RGA5. Plant Cell 29, 156–168

27. Guo, L., Cesari, S., de Guillen, K., Chalvon, V., Mammri, L., Ma, M., Meusnier, I., Bonnot, F., Padilla, A., Peng, Y. L., Liu, J., and Kroj, T. (2018) Specific recognition of two MAX effectors by integrated HMA domains in plant immune receptors involves distinct binding surfaces. Proc Natl Acad Sci U S A 115, 11637–11642

28. Costanzo, S., and Jia, Y. L. (2010) Sequence variation at the rice blast resistance gene Pi-km locus: Implications for the development of allele specific markers. Plant Sci 178, 523–530

29. Kanzaki, H., Yoshida, K., Saitoh, H., Fujisaki, K., Hirabuchi, A., Alaux, L., Fournier, E., Tharreau, D., and Terauchi, R. (2012) Arms race co-evolution of Magnaporthe oryzae AVR-Pik and rice Pik genes driven by their physical interactions. Plant J 72, 894–907

30. Yoshida, K., Saitoh, H., Fujisawa, S., Kanzaki, H., Matsumura, H., Tosa, Y., Chuma, I., Takano, Y., Win, J., Kamoun, S., and Terauchi, R. (2009) Association genetics reveals three novel avirulence genes from the rice blast fungal pathogen Magnaporthe oryzae. Plant Cell 21, 1573–1591

31. Varden, F. A., Saitoh, H., Yoshino, K., Franceschetti, M., Kamoun, S., Terauchi, R., Banfield, M.J. (2019) Cross-reactivity of a rice NLR immune receptor to distinct effectors from the blast pathogen leads to partial disease resistance. bioRxiv

32. Islam, M. T., Croll, D., Gladieux, P., Soanes, D. M., Persoons, A., Bhattacharjee, P., Hossain, M. S., Gupta, D. R., Rahman, M. M., Mahboob, M. G., Cook, N., Salam, M. U., Surovy, M. Z., Sancho, V. B., Maciel, J. L., NhaniJunior, A., Castroagudin, V. L., Reges, J. T., Ceresini, P. C., Ravel, S., Kellner, R., Fournier, E., Tharreau, D., Lebrun, M. H., McDonald, B. A., Stitt, T., Swan, D., Talbot, N. J., Saunders, D. G., Win, J., and Kamoun, S. (2016) Emergence of wheat blast in Bangladesh was caused by a South American lineage of Magnaporthe oryzae. BMC Biol 14, 84

33. Michelmore, R., Coaker, G., Bart, R., Beattie, G., Bent, A., Bruce, T., Cameron, D., Dangl, J., Dinesh-Kumar, S., Edwards, R., Eves-van den Akker, S., Gassmann, W., Greenberg, J. T., Hanley-Bowdoin, L., Harrison, R. J., Harvey, J., He, P., Huffaker, A., Hulbert, S., Innes, R., Jones, J. D. G., Kaloshian, I., Kamoun, S., Katagiri, F., Leach, J., Ma, W., McDowell, J., Medford, J., Meyers, B., Nelson, R., Oliver, R., Qi, Y., Saunders, D., Shaw, M., Smart, C., Subudhi, P., Torrance, L., Tyler, B., Valent, B., and Walsh, J. (2017) Foundational and Translational Research Opportunities to Improve Plant Health. Mol Plant Microbe Interact 30, 515–516

34. Savary, S., Willocquet, L., Pethybridge, S. J., Esker, P., McRoberts, N., and Nelson, A. (2019) The global burden of pathogens and pests on major food crops. Nat Ecol Evol 3, 430–439

35. Bomblies, K., Lempe, J., Epple, P., Warthmann, N., Lanz, C., Dangl, J. L., and Weigel, D. (2007) Autoimmune response as a mechanism for a Dobzhansky-Muller-type incompatibility syndrome in plants. PLoS Biol 5, e236

36. Altpeter, F., Springer, N. M., Bartley, L. E., Blechl, A. E., Brutnell, T. P., Citovsky, V., Conrad, L. J., Gelvin, S. B., Jackson, D. P., Kausch, A. P., Lemaux, P. G., Medford, J. I., Orozco-Cardenas, M. L., Tricoli, D. M., Van Eck, J., Voytas, D. F., Walbot, V., Wang, K., Zhang, Z. J., and Stewart, C. N., Jr. (2016) Advancing Crop Transformation in the Era of Genome Editing. Plant Cell 28, 1510–1520

37. Watson, A., Ghosh, S., Williams, M. J., Cuddy, W. S., Simmonds, J., Rey, M. D., Asyraf Md Hatta, M., Hinchliffe, A., Steed, A., Reynolds, D., Adamski, N. M., Breakspear, A., Korolev, A., Rayner, T., Dixon, L. E., Riaz, A., Martin, W., Ryan, M., Edwards, D., Batley, J., Raman, H., Carter, J., Rogers, C., Domoney, C., Moore, G., Harwood, W., Nicholson, P., Dieters, M. J., DeLacy, I. H., Zhou, J., Uauy, C., Boden, S. A., Park, R. F., Wulff, B. B. H., and Hickey, L. T. (2018) Speed breeding is a powerful tool to accelerate crop research and breeding. Nat Plants 4, 23–29

38. Arora, S., Steuernagel, B., Gaurav, K., Chandramohan, S., Long, Y., Matny, O., Johnson, R., Enk, J., Periyannan, S., Singh, N., Asyraf Md Hatta, M., Athiyannan, N., Cheema, J., Yu, G., Kangara, N., Ghosh, S., Szabo, L. J., Poland, J., Bariana, H., Jones, J. D. G., Bentley, A. R., Ayliffe, M., Olson, E., Xu, S. S., Steffenson, B. J., Lagudah, E., and Wulff, B. B. H. (2019) Resistance gene cloning from a wild crop relative by sequence capture and association genetics. Nat Biotechnol 37, 139–143

39. Berrow, N. S., Alderton, D., Sainsbury, S., Nettleship, J., Assenberg, R., Rahman, N., Stuart, D. I., and Owens, R. J. (2007) A versatile ligation-independent cloning method suitable for high-throughput expression screening applications. Nucleic Acids Res 35, e45

40. Engler, C., Kandzia, R., and Marillonnet, S. (2008) A one pot, one step, precision cloning method with high throughput capability. PLoS One 3, e3647

41. Lobstein, J., Emrich, C. A., Jeans, C., Faulkner, M., Riggs, P., and Berkmen, M. (2012) SHuffle, a novel Escherichia coli protein expression strain capable of correctly folding disulfide bonded proteins in its cytoplasm. Microb Cell Fact 11, 56

42. Studier, F. W. (2005) Protein production by auto-induction in high density shaking cultures. Protein Expr Purif 41, 207–234

43. Myszka, D. G. (1999) Improving biosensor analysis. J Mol Recognit 12, 279–284

44. Winter, G. (2010) xia2: an expert system for macromolecular crystallography data reduction. Journal of Applied Crystallography 43, 186–190

45. Winn, M. D., Ballard, C. C., Cowtan, K. D., Dodson, E. J., Emsley, P., Evans, P. R., Keegan, R. M., Krissinel, E. B., Leslie, A. G., McCoy, A., McNicholas, S. J., Murshudov, G. N., Pannu, N. S., Potterton, E. A., Powell, H. R., Read, R. J., Vagin, A., and Wilson, K. S. (2011) Overview of the CCP4 suite and current developments. Acta Crystallogr D Biol Crystallogr 67, 235–242

46. McCoy, A. J., Grosse-Kunstleve, R. W., Adams, P. D., Winn, M. D., Storoni, L. C., and Read, R. J. (2007) Phaser crystallographic software. J Appl Crystallogr 40, 658–674

47. Emsley, P., Lohkamp, B., Scott, W. G., and Cowtan, K. (2010) Features and development of Coot. Acta Crystallogr D Biol Crystallogr 66, 486–501

48. Murshudov, G. N., Skubak, P., Lebedev, A. A., Pannu, N. S., Steiner, R. A., Nicholls, R. A., Winn, M. D., Long, F., and Vagin, A. A. (2011) REFMAC5 for the refinement of macromolecular crystal structures. Acta Crystallogr D Biol Crystallogr 67, 355–367

49. Chen, V. B., Arendall, W. B., 3rd, Headd, J. J., Keedy, D. A., Immormino, R. M., Kapral, G. J., Murray, L. W., Richardson, J. S., and Richardson, D. C. (2010) MolProbity: all-atom structure validation for macromolecular crystallography. Acta Crystallogr D Biol Crystallogr 66, 12–21

